# Elevated trace elements in *Posidonia oceanica* and *Cymodocea nodosa* at six Mediterranean volcanic seeps

**DOI:** 10.1101/433987

**Authors:** A.K. Mishra, R. Santos, J.M. Hall-Spencer

## Abstract

Seagrasses form important habitats around shallow marine CO_2_ seeps, providing opportunities to assess trace element (TE) accumulation along gradients in seawater pH. Here we assessed Cd, Cu, Hg, Ni, Pb and Zn levels in sediment and seagrasses at six CO_2_ seeps and reference sites off Italy and Greece. Some seep sediments had much higher concentrations of TEs, the extreme example being Cd at 43-fold above reference levels. Sediment Quality Guideline (SQG) scores indicated that three seeps had sediment TEs levels likely to have “Adverse impacts” on marine biota; namely Vulcano (for Hg), Ischia (for Cu) and Paleochori (for Cd and Ni). SQG indicated seep sediments of Italian seeps were adversely affected by Cu and Hg, whereas Greek CO_2_ seeps were affected by Cd and Ni. An increase in sediment TEs levels positively corelated with higher levels of TEs in seagrass roots of *Posidonia oceanica* (Zn and Ni) and *Cymodocea nodosa* (Zn). Differences in the bioavailability and possible toxicity of TEs helps explain why seagrasses were abundant at some CO_2_ seeps but not others.

## Introduction

Around 30% of anthropogenic CO_2_ emissions dissolve into the surface ocean causing the pH to fall and altering seawater carbonate chemistry leading to ocean acidification (OA) (Caldeira and Wickett, 2003). These chemical changes are a major threat to marine species and ecosystems, the reasons why the United Nations Sustainable Development Goal 14.3 is to “Minimize and address the impacts of ocean acidification” (United Nations, 2015). Rising CO_2_ levels are expected to reduce marine biodiversity and alter trophic interactions (Kroeker et al., 2013; Sunday et al., 2016) which will impact a range of ecosystem services (Lemasson et al., 2017).

Metallic trace elements (both essential and non-essential) in marine ecosystem occur naturally in very low concentrations (Alloway, 1995) and are non-toxic. However, at sufficient high enough concentration elements such as As, Cu, Pb and Hg can be toxic and harmful to the coastal biota (Stumm Morgan, 1995). The toxicity of these elements depends on their chemical form, As for example is toxic when found in its metalloid form, Hg and Pb are toxic as free ions and Cu is toxic when reduced to Cu (I) (Tchounwou et al., 2014). A major concern is that OA will exacerbate the harmful effects of metal pollution, which is a widespread problem in coastal ecosystems (Ivanina et al., 2015; Lewis et al., 2016). OA is expected to increase the bioavailability and toxicity of metals both in sediments (Roberts et al., 2013) and the water column (Millero et at., 2009). Lower pH combined with low oxygen levels can release metals to water column that are bound to sediments (Atkinson et al., 2007) and low pH can also alter the speciation of elements like Cu, Ni and Zn (Zeng et al., 2015) resulting in increased toxicity. However, the level of toxicity on marine organisms will depend on the uptake rate and interaction of metals at the receptor sites of an organism (Batley et al., 2004). Studies indicate that, the uptake and availability of elements like Cd, Co, Cu, Hg, Ni, Pb and Zn are going to increase under OA and low pH range (8.1 to 7.8) predicted for 21^st^ century (Byrne et al., 1998; Richards et al., 2011). For example, seawater free ion concentration of Cu is expected to increase by 115% (Pascal et al., 2010; Richards et al., 2011), Pb by 4.56% (Millero et al., 2009; Dong et al., 2016) whereas Cd concentration may decrease or be unaffected (Pascal et al., 2010) under OA and low pH scenarios. But the increase in free ions is predicted to increase the toxicity of metals under OA (Lacoue-Labarthe et al., 2009; 2012).

Most studies on the bioavailability of trace metals at elevated CO_2_ have been carried out in laboratory conditions (Besar et al., 2008; Richir & Gobert, 2013; Bravo et al., 2016) which undermines the complex behaviour of trace elements in marine environment. The ecological risk posed by the effects of OA on metal contaminated water column and sediments is difficult to assess in the laboratory (Millero et al., 2009), which emphasizes the importance of studies on the interplay between trace elements and OA in-situ. Submarine hydrothermal activity is of interest as this causes natural gradients in both ocean acidity and trace elements, providing natural conditions to assess their combined effects (Monia Renzi et al., 2011; Kadar et al., 2012; Vizzini et al., 2013) on marine biota. While relationships among organisms, environmental factors and trace elements have received much attentions at deep sea hydrothermal vents (Kadar et al., 2007; Cravo et al., 2007), those of shallow marine CO_2_ seeps are still little understood.

Shallow water CO_2_ seeps represent natural analogues for future coastal ecosystems, providing entire seabed of marine communities exposed to the shifts in carbonate chemistry (low pH) expected for 21^st^ century (Hall-Spencer et al., 2008; Enochs et al., 2015; Connell et al., 2017). At such seeps, there are often elevated levels of trace elements and hydrogen sulphide (H_2_S) (Kadar et al., 2012; Boatta et al., 2013; Vizzini et al., 2013) combined with low pH that may be harmful to marine biota (Barry et al., 2010; Lauritano et al., 2015; Olivé et al., 2017). At these CO_2_ seeps, habitat forming marine fauna (such as macroalgae and seagrass) are observed (Apostolaki et al., 2014; Vizzini et al., 2010) which provides a window of opportunity to investigate the combined effects of OA and metal levels on seagrass. However, the seagrass population are not evenly abundant around these CO_2_ seeps, for example lush stands of seagrass are observed at some seeps off Ischia in the Mediterranean Sea and at several sites around Papua New Guinea (Hall-Spencer et al., 2008; Russel et al., 2013), whereas seagrass abundance is less at CO_2_ seeps off Panarea and Vulcano off Italy, where H_2_S and elevated trace elements appear to affect the plants (Vizzini et al., 2010, 2013). Saying that, studies on seagrass eco-physiology at these CO_2_ seeps have provided mixed results, for example, *Posidonia oceanica* antioxidant stress related genes are more expressed near volcanic seeps (at Panarea and Ischia Islands, Italy), whereas the expression of genes involved in photosynthesis and growth responses of *Cymodocea nodosa* near a seep off Vulcano islands, Italy were contrary to the expected beneficial effects of CO_2_ (Olivé et al., 2017). On the other hand, under experimental CO_2_ enrichment there was a significant increased expression of *C. nodosa* transcripts associated with photosynthesis, as expected (Ruocco et al., 2017).

Seagrass are important coastal habitats due to their high productivity and biodiversity (Thom, 2001). They provide food nurseries for fish, turtles and mammals (Coles et al., 2007), they can be major carbon sinks and can sequester contaminants such as excess nutrients and metals (Orth et al., 2006; Fourqurean et al., 2012). Seagrass productivity is predicted to increase as CO_2_ levels continue to rise if temperature increases do not become too stressful (Koch et al., 2013; Brodie et al., 2014). Seagrasses accumulate trace elements and so are used as bioindicator in coastal ecosystems (Catsiki and Panayotidis, 1993). The plants take in trace elements via the roots, rhizomes or the leaves and can translocate them between these tissue compartments (Ralph et al., 2006). The rate of uptake varies between essential and non-essential trace elements and between tissues and this introduces these substances into the food web via grazing and decomposition (Lewis and Devereux, 2009). For instance, *Cymodocea serrulata* transfers trace elements to detrital feeders (Klumpp et al., 1989).

Laboratory studies have shown that, at elevated CO_2_ and above threshold levels Cu, Pb and Zn exert toxic effects on the physiology of the seagrasses *Zostera capricorni* (Ambo-Rappe et al., 2007) and *Halophila ovalis* (Ambo-Rappe et al., 2011). Many CO_2_ seeps around Greece and Italy have seagrass (Hall-Spencer et al., 2008; Vizzini et al., 2010; Apostolaki et al., 2014). Little is known about the trace metal content, their accumulation in seagrass roots, rhizomes and leaves at CO_2_ seeps..

Here we expand on the work undertaken by Vizzini et al., (2013) to quantify the concentrations of trace elements, in sediments and seagrass at multiple seep sites around the Mediterranean. Our aim was to find out whether increased levels of trace elements near seeps correlate with increases in trace elements accumulation in seagrass roots, rhizomes and leaves and whether seagrass have preferences in metal accumulation patterns.

## Methods

### Study sites

We surveyed six sites in the Mediterranean Sea, all of which had seagrasses (*Posidonia oceanica* or *Cymodocea nodosa*) growing on sandy bottom in high salinity and high alkalinity (Table 1). At each site, a high CO_2_ station and a reference station were sampled between May - July 2014. The annual temperature range was around 18-22°C for all six locations and the CO_2_ seeps were at 0-10 m depth with a tidal range of 0.30-0.50 m.

**Table 1:**
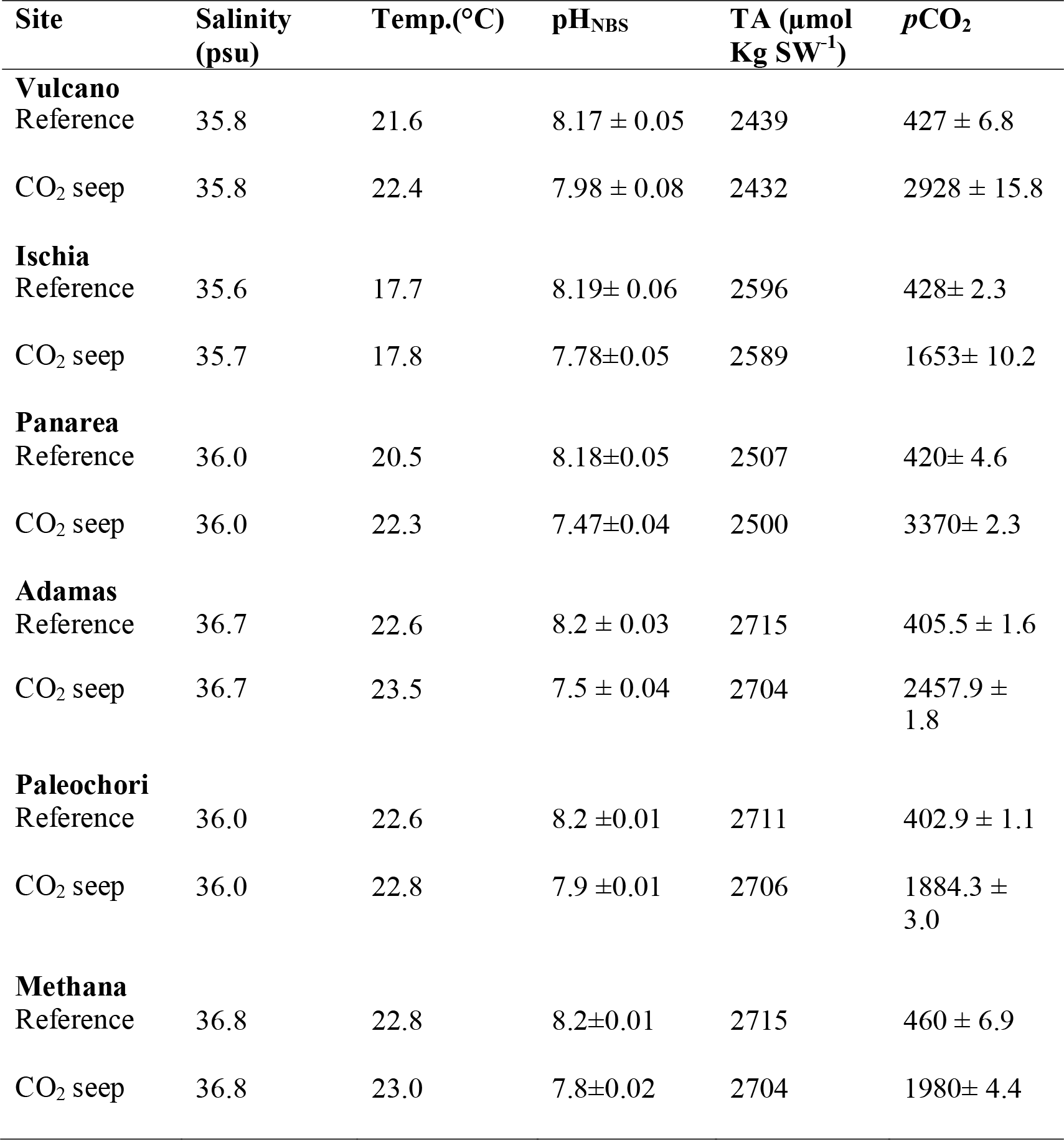
Seawater salinity, temperature, total alkalinity, pH and *p*CO_2_ values (mean ± SE, n=5) at six Mediterranean CO_2_ seeps and reference sites between May-July 2014.

| Site | Salinity (psu) | Temp.(°C) | pH<sub>NBS</sub> | TA (μmol Kg SW<sup>-1</sup>) | pCO<sub>2</sub> |
| --- | --- | --- | --- | --- | --- |
| **Vulcano** |  |  |  |  |  |
| Reference | 35.8 | 21.6 | 8.17 ± 0.05 | 2439 | 427 ± 6.8 |
| CO<sub>2</sub> seep | 35.8 | 22.4 | 7.98 ± 0.08 | 2432 | 2928 ± 15.8 |
| **Ischia** |  |  |  |  |  |
| Reference | 35.6 | 17.7 | 8.19± 0.06 | 2596 | 428± 2.3 |
| CO<sub>2</sub> seep | 35.7 | 17.8 | 7.78±0.05 | 2589 | 1653± 10.2 |
| **Panarea** |  |  |  |  |  |
| Reference | 36.0 | 20.5 | 8.18±0.05 | 2507 | 420± 4.6 |
| CO<sub>2</sub> seep | 36.0 | 22.3 | 7.47±0.04 | 2500 | 3370± 2.3 |
| **Adamas** |  |  |  |  |  |
| Reference | 36.7 | 22.6 | 8.2 ± 0.03 | 2715 | 405.5 ± 1.6 |
| CO<sub>2</sub> seep | 36.7 | 23.5 | 7.5 ± 0.04 | 2704 | 2457.9 ± 1.8 |
| **Paleochori** |  |  |  |  |  |
| Reference | 36.0 | 22.6 | 8.2 ±0.01 | 2711 | 402.9 ± 1.1 |
| CO<sub>2</sub> seep | 36.0 | 22.8 | 7.9 ±0.01 | 2706 | 1884.3 ± 3.0 |
| **Methana** |  |  |  |  |  |
| Reference | 36.8 | 22.8 | 8.2±0.01 | 2715 | 460 ± 6.9 |
| CO<sub>2</sub> seep | 36.8 | 23.0 | 7.8±0.02 | 2704 | 1980± 4.4 |

### Vulcano, Italy

We sampled Levante Bay (38.4 N, 15.0 E) off Vulcano island (Fig. 1A). The underwater gas emissions are 97-98% CO_2_ with 2.2% hydrogen sulfide (H_2_S) close to the seeps, decreasing to less than 0.005% H_2_S towards the north-eastern part of the bay (Capaccioni et al., 2001; Milazzo et al., 2014). *Cymodocea nodosa* was absent near the main vents so we, collected it on the periphery of the CO_2_ seeps at 1 m depth.

**Fig. 1.**
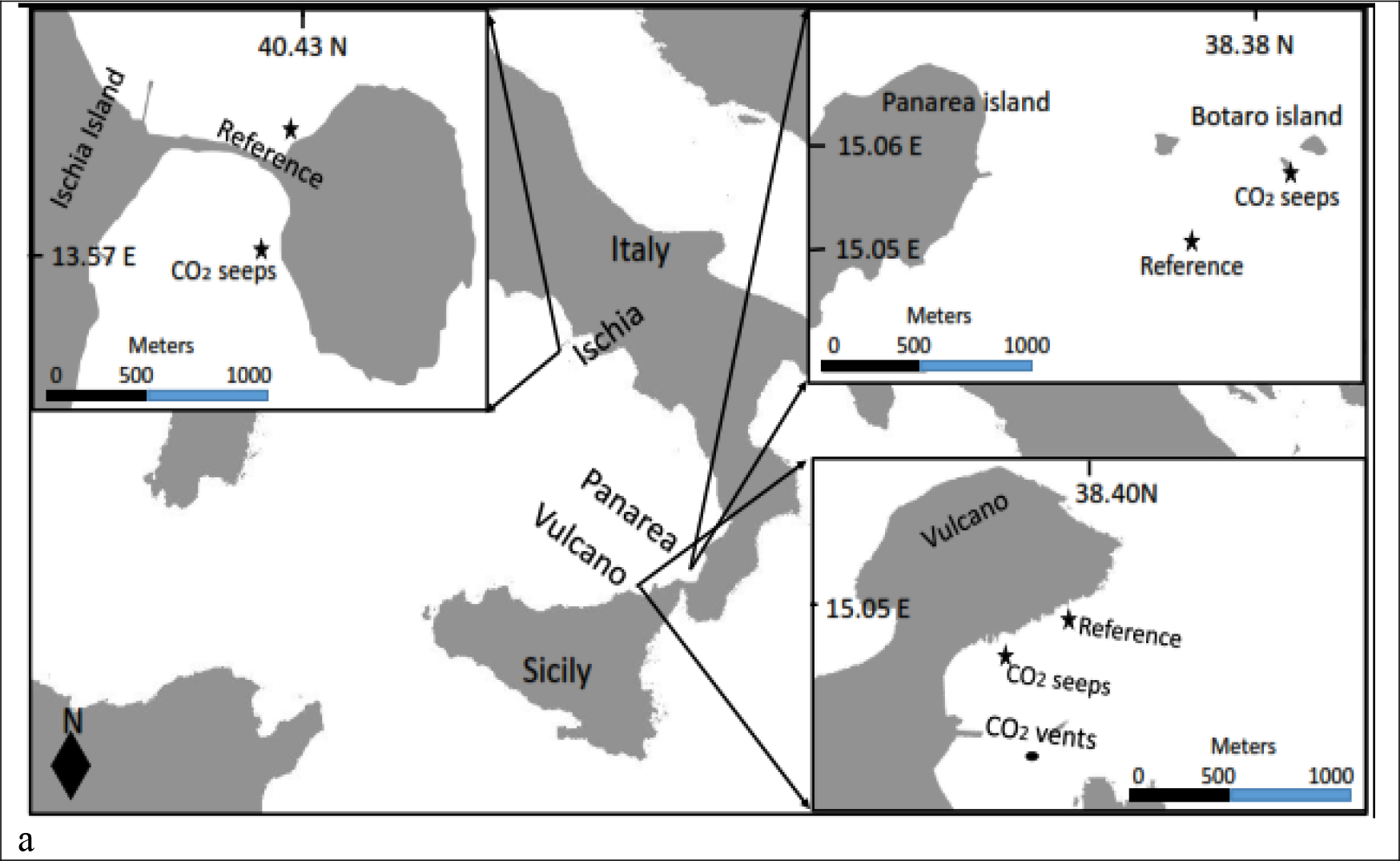
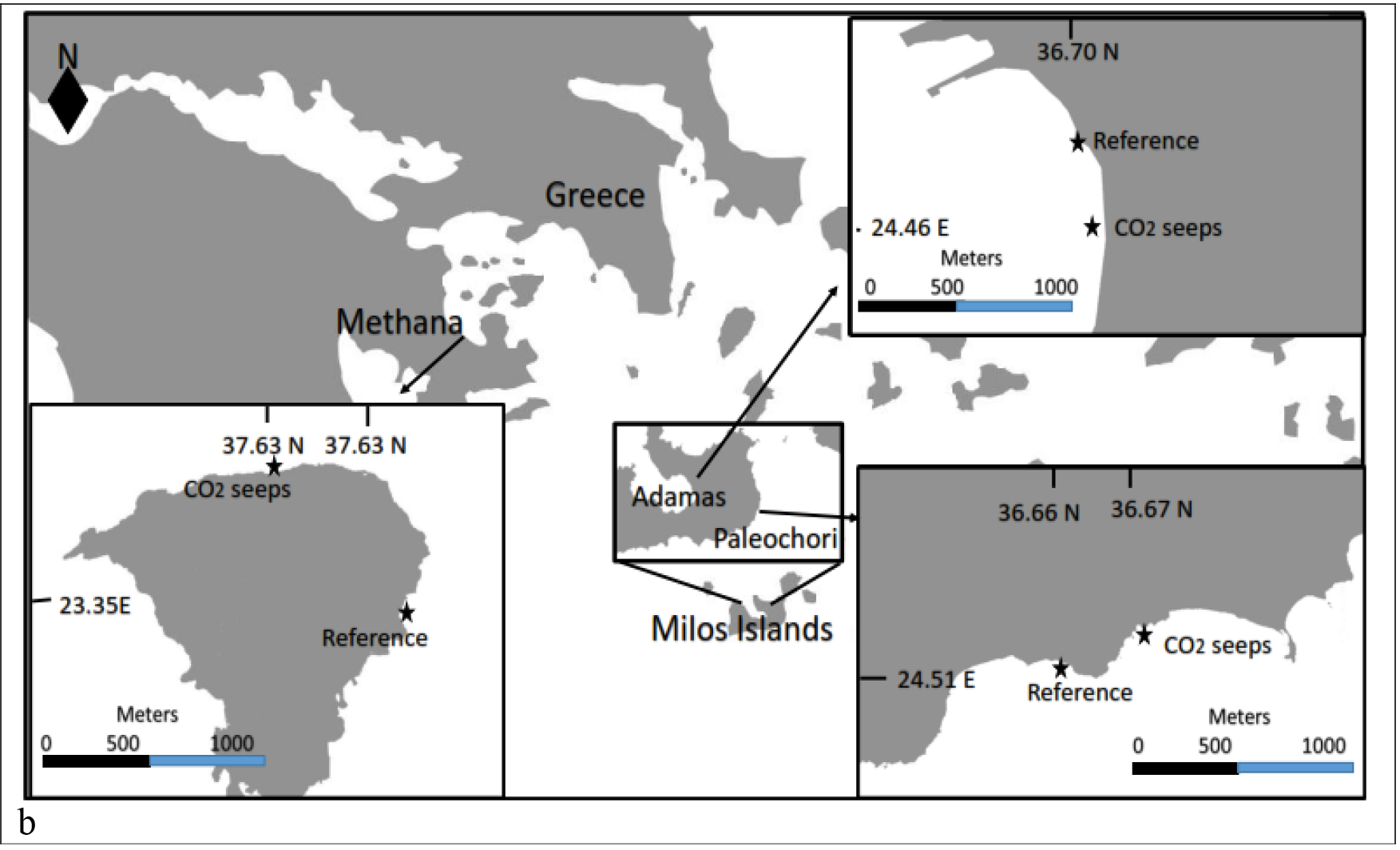
Study sites in Italy a) and b) Greece, showing reference and CO_2_ seep sites, which were all sampled between May to July 2014.

### Ischia, Italy

At the Castello Aragonese, off Ischia (40°43’50.4”N; 13°57’48.2”E) CO_2_ bubbles up in shallow water seeps (Fig. 1A). Here the gas is 90–95% CO_2_, 3–6% N_2_, 0.6–0.8% O_2_, 0.2–0.8% CH_4_ and 0.08–0.1% air and the seeps lack H_2_S (Tedesco, 1996). Here *Posidonia oceanica* meadows were sampled at 0.5m depth from the seep area and from a northward reference site with no CO_2_ bubbling (Fig.2a).

**Fig. 2.**
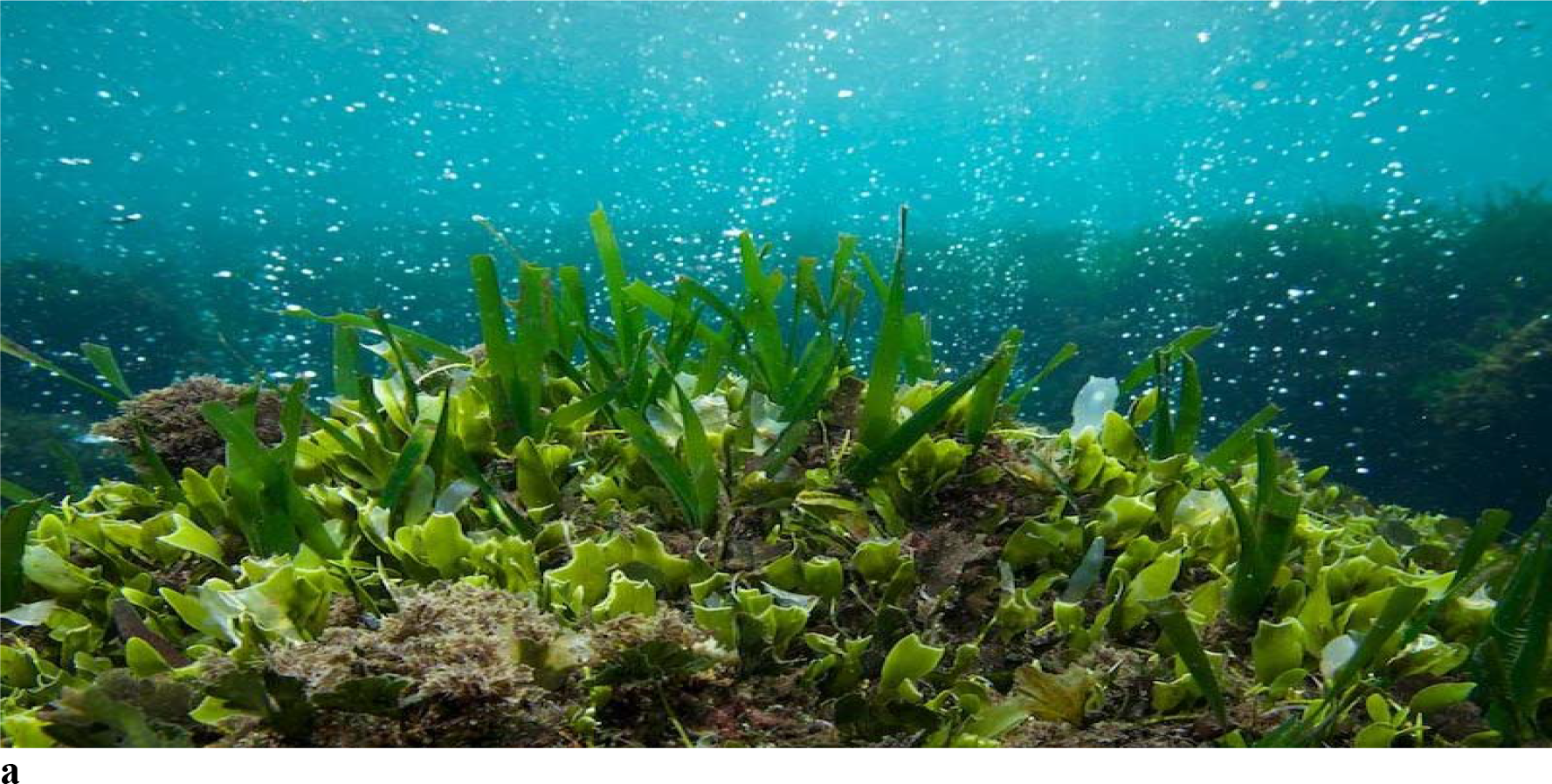
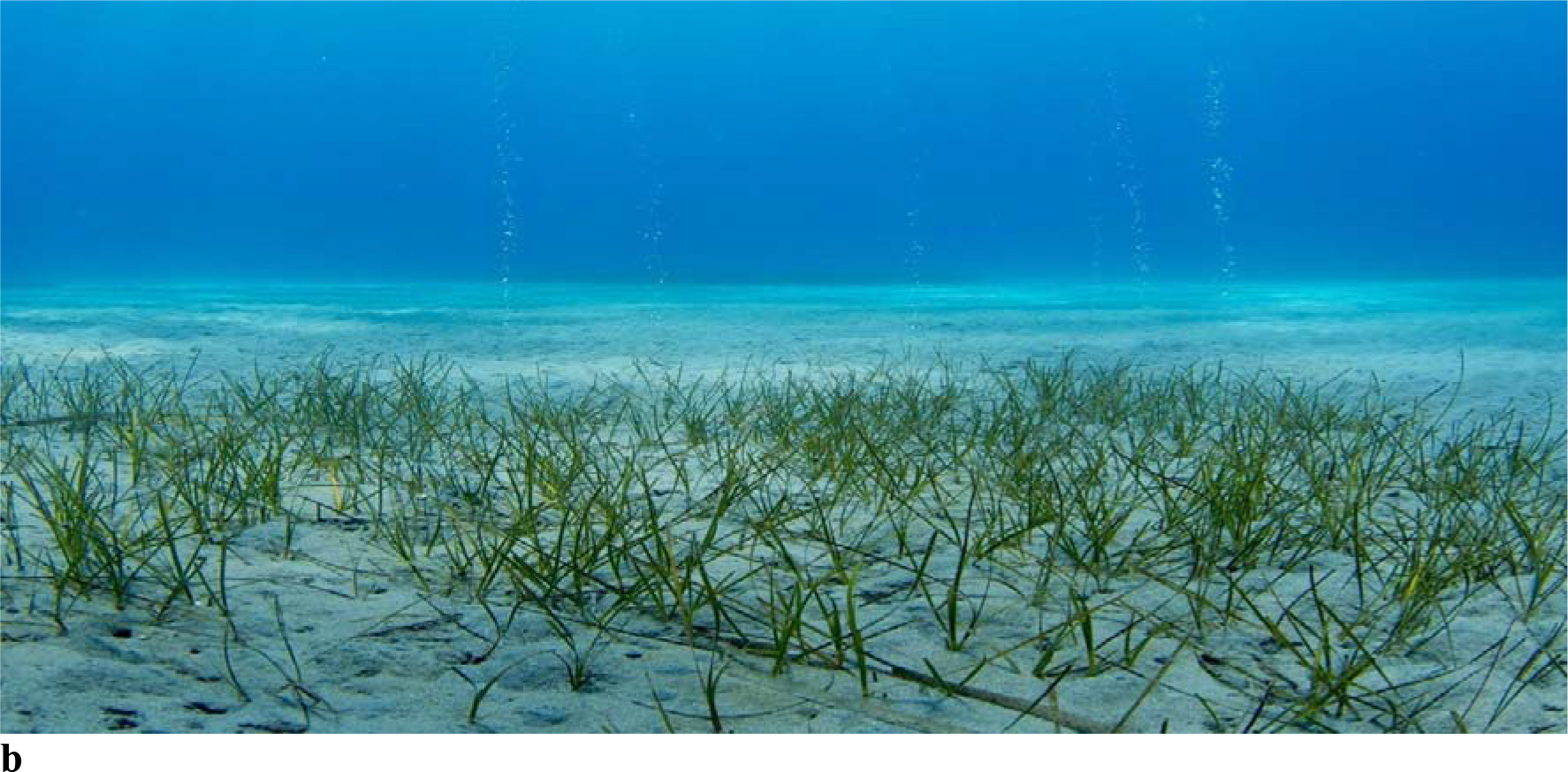
a) *Posidonia oceanica* and b) *Cymodocea nodosa* meadows at CO_2_ seeps off Ischia (Italy) and Paleochori (Greece). Photo credits for a) *Posidonia oceanica*, and b) *Cymodocea nodosa* meadows at Italy and Greece: Jason Hall Spencer, University of Plymouth, UK and Thanos Dailianis of Hellenic Centre for Marine Research, Greece respectively.

### Panarea, Italy

Panarea island (38°38’12.2”N; 15°06’42.5”E) is part of the Aeolian Archipelago in the Southern Tyrrhenian Sea (Fig.1A). On the main island and on the surrounding seafloor, tectonic faults have many gas seeps (Gabianelli et al., 1990; Voltattorni et al., 2009). The underwater gas emissions around these seeps are 92-95%CO_2_, 2.99-6.23% N_2_, 0.69-1.2% O_2_ and 0.65-3% H_2_S (Caramanna et al., 2010). Here *P. oceanica* was sampled at 5m depth.

### Milos Islands, Greece

Adamas thermal springs (36.70 N, 24.46 E) and Paleochori Bay (36.67 N, 24.51 E) are situated on southwest and southeast part of Milos island respectively (Fig.1B). Milos island is part of an extensive submarine venting, from the intertidal to depths of more than 100 m (Dando et al.,1999). The underwater gas seeps at Adamas and Paleochori are located <1 to 4 m depth. The released gases are mainly composed of 92.5% CO_2_ with some CH_4_ and H_2_ (Bayraktarov et al., 2013). *Cymodocea nodosa* meadows were sampled at 4m depth at Paleochori Bay and 2m at Adamas thermal station (Fig.2b).

### Methana, Greece

The Methana peninsula (37.638428 N; 23.359730 E) is the westernmost volcanic system of the northern Aegean Volcanic Arc (Fig.1B), derived from the subduction of the African tectonic plate beneath the Eurasian plate. We sampled the area described by Baggini et al., (2014) near Agios Nikolaos village on the NE part of the peninsula. The gases were 90% CO_2_, with small amounts of nitrogen, carbon monoxide and methane (D’Alessandro *et al.*, 2008). Here we sampled *Posidonia oceanica* meadows at 8-10 m depth.

### Water sampling

Water samples (n=5) were collected at each CO_2_ seep and Reference station of all six sites in 100 ml Winkler bottles and were fixed with 20 μl mercuric chloride in the field, stored in dark cool- boxes and transported to the laboratory for total alkalinity (TA) analysis. The pH_NBS_ (using pH meter, Titrino Methron, Thermo Scientific) and temperature of the water samples were measured in the field immediately after collection. In the laboratory 80 ml water samples were analysed for TA using a Lab Titrino analyser following methods given by Dickson et al., (2007). Sterilized sea water was used as reference materials (CRM Batch 129, accuracy-98.7% Dickson, 2013) for TA analysis. Hanna buffer (HI7007L, Hanna Instruments, accuracy-99%) was used to calibrate the pH electrodes. Temperature, pH_NBS_ and TA data were used to calculate *p*CO_2_ using CO_2_SyS program following methods given by Pierrot et al., (2006). Dissociation constants (K_1_ and K_2_) developed by Meherbach et al., (1973) and refitted by Dickson et al., (1987) and dissociated constant for boric acid (K_B_) developed by Dickson et al., (2007) was used in *p*CO_2_ calculation.

### Sediment & seagrass sampling

Sediment samples (n=5) mostly sand was collected from CO_2_ seep and Reference stations of all six sites by SCUBA diving. A 10-cm long and 2 cm diameter syringe with the tip cut off to was used to suck up the upper 5 cm of sand. From each station (CO_2_ seep and Reference) five points were selected randomly (1m apart) to collect samples. The sediment samples were stored in plastic bags in dark boxes and transferred to lab. In the laboratory they were dried at 40°C in an oven till a constant weight was achieved and then analysed for the grain size following dry sieving at Half Phi intervals (Blott and Pye, 2001). After grain size analysis the fine and very fine of sediment fraction (<180-63 μm) were collected and stored in plastic bottles for trace metal analysis.

Samples (n=5, whole plants) of *Cymodocea nodosa* (from Vulcano, Adamas and Paleochori islands) and *Posidonia oceanica* (from Ischia, Panarea and Methana) were collected by SCUBA diving up to 10-m depth at the CO_2_ seep and Reference stations of all six sites. The seagrasses were rinsed well to remove sediments, razor scraped to remove leaf epiphytes, leaf scales removed from rhizomes (*P. oceanica*) by hand and with soft toothbrush and then washed with distilled water, air-dried and stored in polybags until analyses. Seagrass leaves, roots and rhizomes were oven dried at 40°C and powdered in a mortar and stored till further analysis.

### Analytical Methods

Total trace elements (Cd, Cu, Hg, Ni, Pb and Zn) concentration was determined using Aqua Regia Soluble Total method (Modified by Laboratory of the Government Chemist (LGC) UK from ISO11466). Dried sediment (0.25 g) was put into digestion tubes (Tecator type). Cold and concentrated acids in the order: 4.5 mL Hydrochloric acid (HCl,): 1.5 mL Nitric acid (HNO_3_) was added to the tubes. The digestion tubes were left to pre-digest, for one hour then heated for 2 hours at 95 - 100°C. After cooling, the digest was filtered quantitatively into a volumetric flask and diluted using 2% HNO_3_ (25 ml volume).

For dried seagrass (leaves, rhizomes and roots), 0.25g of sample was added to 6mL of HNO_3_ following the same procedure as metals and the volume was made up to 25mL. Similarly, blanks and standards (LGC Reference Materials, UK, recovery-95%) used for sediments (LCG6156) and plants (LGC7162) were prepared using the same method. Analysis of Cd, Co, Cu, Hg, Pb and Zn was performed using an ICP-MS (Thermo Scientific, iCAP 7000 Series) and an ICP-AES (Thermo Scientific, X Series-2) in triplicate with analytical detection precision of 99.5%.

All acids were analytical grade. Normal precautions for metal analysis were observed throughout the analytical procedures. HCL (37%w/w) and HNO_3_ (69% w/w) were Ultrapure type (Ultrapure, Fischer Chemicals, USA). All glassware was soaked overnight in 10% HNO_3_ and washed with distilled water and oven dried before use.

### Data Analysis

To assess the sediment quality of all six locations we used Sediment Quality Guidelines Quotient (SQG-Q, Long and MacDonald, 1998). Among the environmental quality indices in the literature, this was chosen for its simplicity, comparability and robustness as reported by Caeiro et al., (2005). The SQG-Q consists of two values: a threshold effects level (TEL) and a probable effect level (PEL) (MacDonald et al., 1996). TEL and PEL, represent concentrations below which adverse biological effects occur rarely and frequently.

The SQG-Q was calculated as follows:

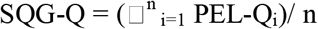

Where PEL-Q_i_ = contaminant/PEL. The PEL-Qi represents the probable effect level quotient (PEL-Q) of the i contaminant and n represents the total number of contaminants (trace metals). Using the SQG-Q index, the sediments were divided into three categories as established by MacDonald et al. (2000). SGQ-Q ≤ 0.1- low potential for adverse biological effects; 0.1< SQG-Q<1- moderate potential for adverse biological effects; SQG-Q≥1- high potential for adverse biological effects.

To assess bio-accumulation of elements, we calculated the Bio Sediment Accumulation Factor (BSAF), which is defined as the ratio between metal concentration in the organism and that in the sediment (Lau et al., 1998; Szefer et al., 1999), given by:

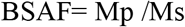

Where Mp is the concentration of the element in the seagrass and Ms is the concentration of the element in the sediment (Fergusson. 1990). BSAF is a key factor in expressing the efficiency of seagrass species to absorb elements from sediments and concentrate specific element in its roots, rhizomes or leaves. Higher BSAF values (>1) indicate a greater capability of accumulation (EPA, 2007).

### Statistics

A three-way ANOVA was used to test for significant differences in trace element concentration among plant compartments (leaves, rhizomes, roots), sediments and stations (Reference, CO_2_ seeps) in each *Posidonia oceanica* sites (Ischia, Panarea and Methana) and each *Cymodocea nodosa* sites (Adamas, Paleochori and Vulcano). All data was pre-checked for normality and homogeneity of variances. When variances were not homogenous, data were ln(x+1) transformed. In some cases, ANOVA main effect was difficult to interpret due to the presence of statistically significant interactions, then Holm-Sidak test was performed for a *posteriori* comparison among levels to check significant main effects in ANOVA. Pearson’s correlation co-efficient was applied to identify correlations between trace element concentration in sediment and seagrass compartments, after testing for normality of distribution on raw or log transformed data. When normality was not achieved, non-parametric Spearman’s rank correlation coefficient was applied. All statistical tests were conducted with a significance level of α =0.05 and data were reported as mean ± standard error (SE).

## Results

Dissolved CO_2_ concentrations were highest (and pH lowest) at each of the seeps; reference sites had normal CO_2_ and pH levels. Among all six CO_2_ seep sites, Panarea was observed with lowest pH (7.47±0.04) level, whereas Vulcano CO_2_ seeps were observed with highest pH (7.98 ± 0.08). The salinity, temperature and total alkalinity were not affected by the seeps (Table 1).

Trace element levels were generally significantly higher in the sediments of seeps than at reference sites (Figs. 3 and 4). The highest differences were found for Ni at Panarea (5.3-fold), Cd at Paleochori (42.6-fold) and Cu at Adamas (8.9- fold) seep sediments than reference sites. Mercury was only observed at CO_2_ seeps off Italy, with the highest level observed within sediment of Vulcano (0.83± 0.15 mg/Kg). Zinc sediment concentrations were similar at all locations but were lowest at Methana (15.67 ± 1.5 mg/Kg). However, Zn levels at the seeps of Panarea were 2.3-fold higher than reference sites.

**Fig. 3.**
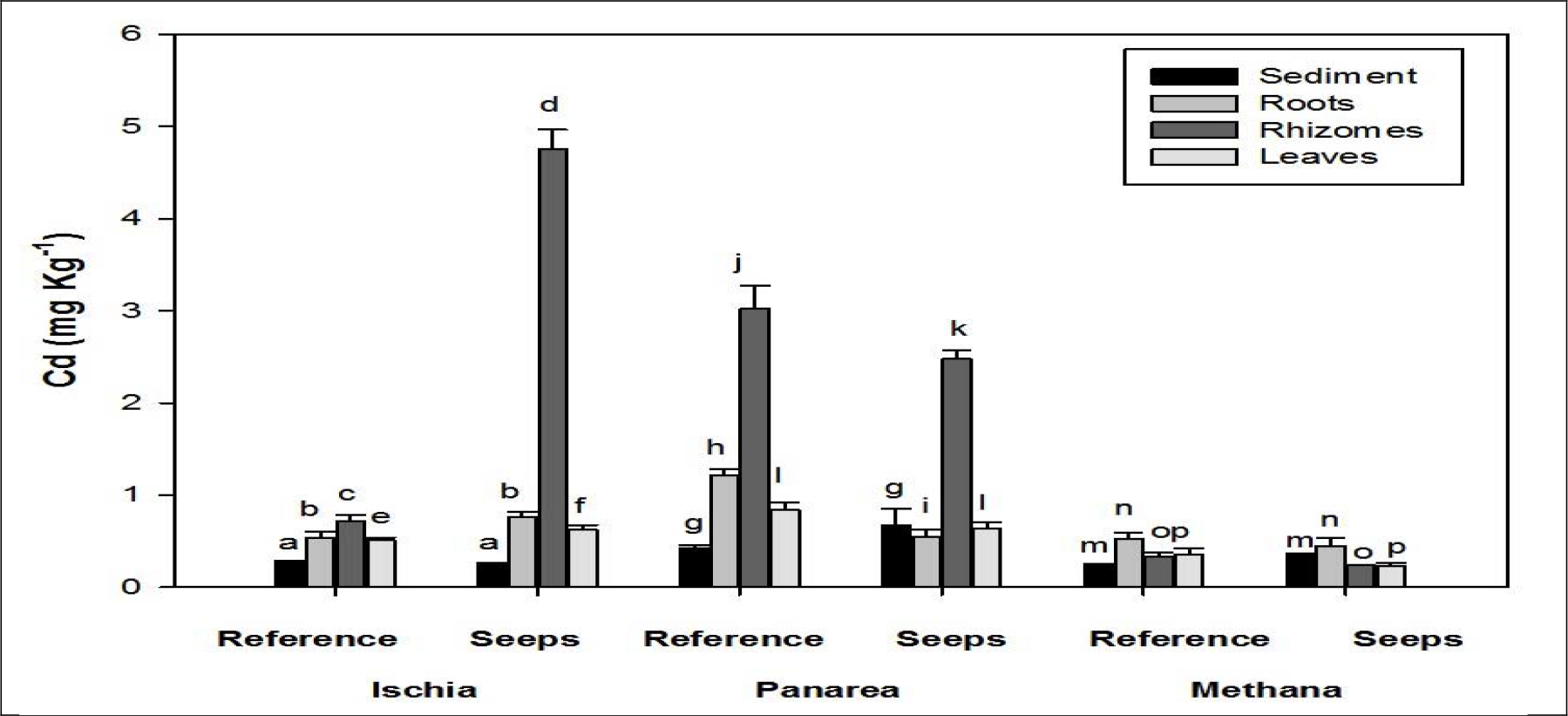
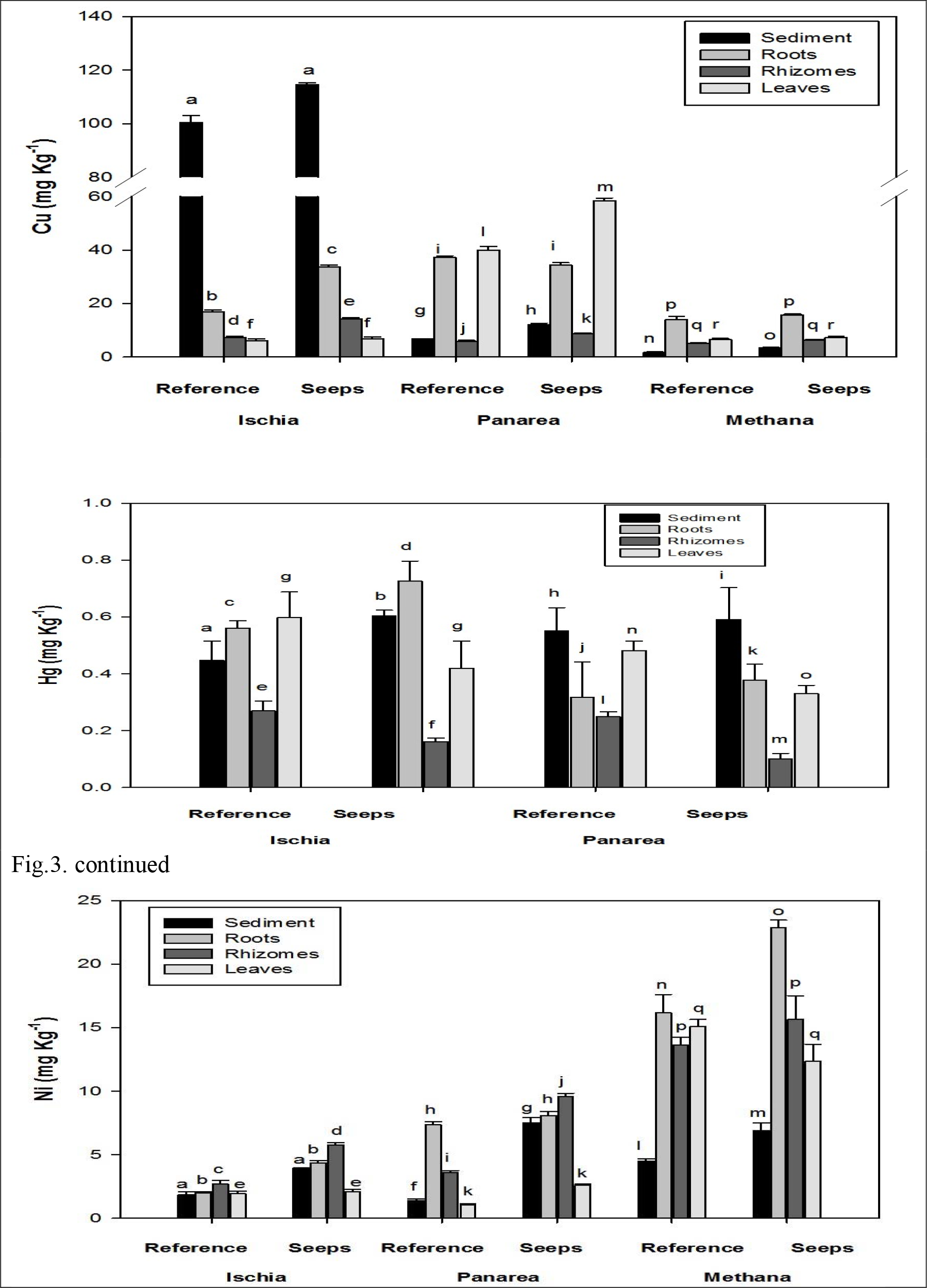
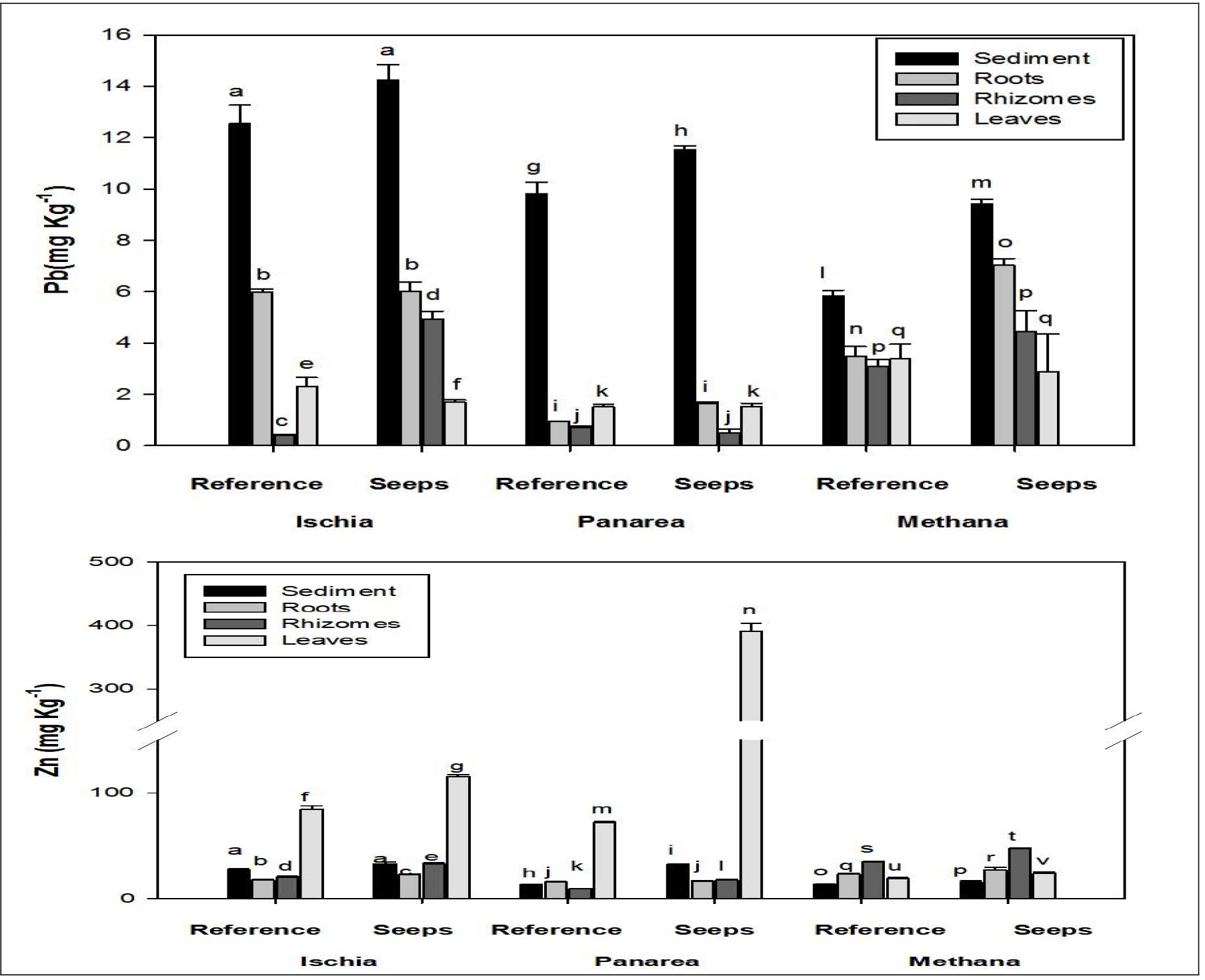
Element concentrations (mean ± SE, n=5) of Cd, Cu, Hg, Ni, Pb and Zn in *Posidonia oceanica* plant compartments and sediments at reference and CO_2_ seep sites off Italy and Greece. Different letters indicate significant differences between reference and CO_2_ seeps site at each location.

Grain size analysis showed that 99% of the sediment particles sampled at all locations were sand(Annex-1). The quality of the seep sediments derived from SQG-Q was usually in the moderate to adverse range than the quality of reference sites at all locations (Table 2). Saying that, the probable ecological risk of trace element levels at CO_2_ seeps were higher with elements such as Hg at Vulcano, Cu at Ischia plus Ni and Cd at Paleochori to have adverse biological effects on seagrass sediment associated biota. (Table 2).

**Table 2.**
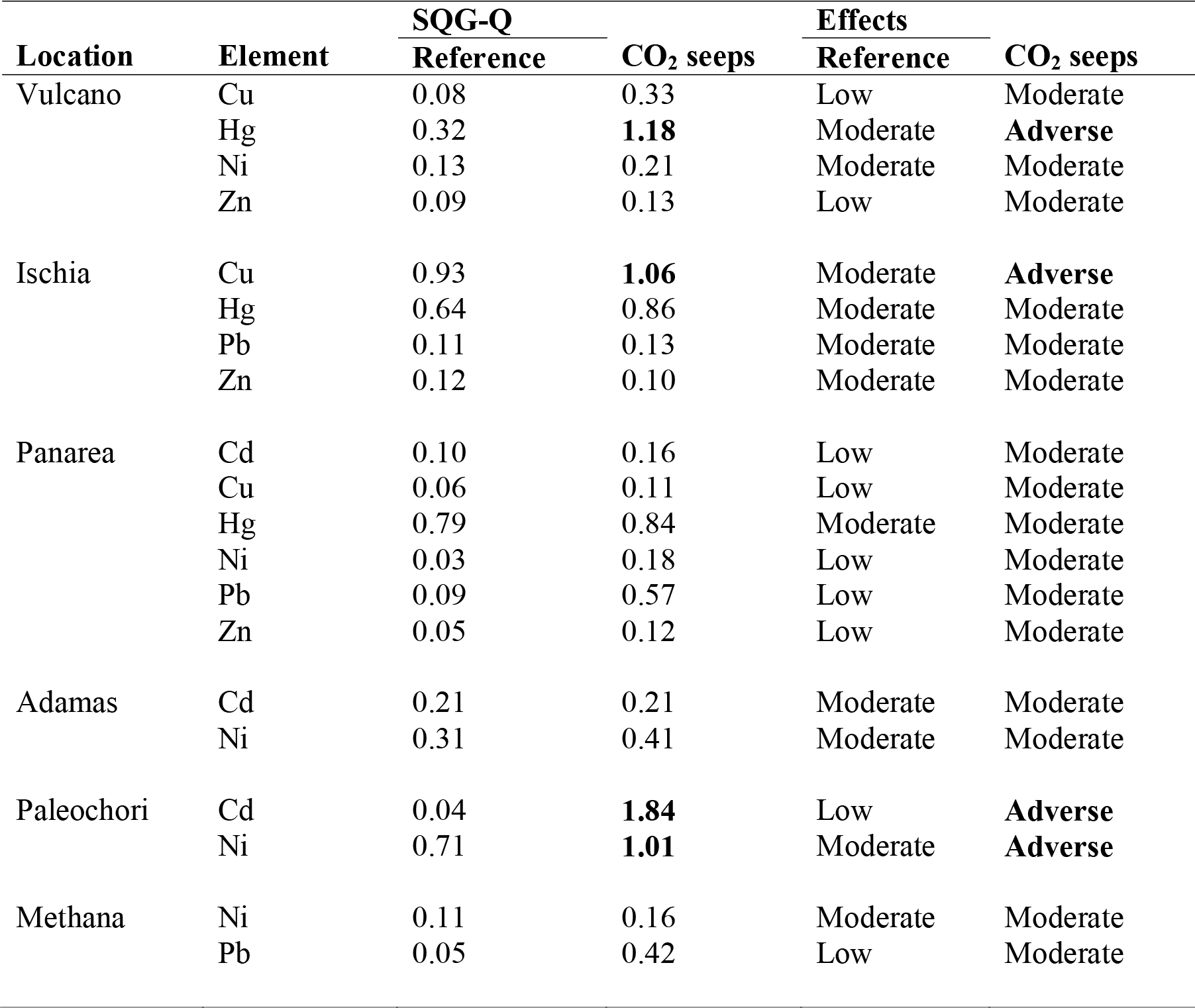
Sediment Quality Guidelines-quotient (SQG-Q) of sediment calculated with Probable Effects Level for Reference and CO_2_ seep sites in Greece and Italy. SQG-Q <0.1 (low effect), <0.1 SQG-Q>1 (moderate effect), SQG-Q>1 (adverse biological effects). Numbers in bold indicate possible adverse effects of trace elements.

| Location | Element | SQG-Q |  | Effects |  |
| --- | --- | --- | --- | --- | --- |
|  |  | Reference | CO<sub>2</sub> seeps | Reference | CO<sub>2</sub> seeps |
| Vulcano | Cu | 0.08 | 0.33 | Low | Moderate |
|  | Hg | 0.32 | **1.18** | Moderate | **Adverse** |
|  | Ni | 0.13 | 0.21 | Moderate | Moderate |
|  | Zn | 0.09 | 0.13 | Low | Moderate |
| Ischia | Cu | 0.93 | **1.06** | Moderate | **Adverse** |
|  | Hg | 0.64 | 0.86 | Moderate | Moderate |
|  | Pb | 0.11 | 0.13 | Moderate | Moderate |
|  | Zn | 0.12 | 0.10 | Moderate | Moderate |
| Panarea | Cd | 0.10 | 0.16 | Low | Moderate |
|  | Cu | 0.06 | 0.11 | Low | Moderate |
|  | Hg | 0.79 | 0.84 | Moderate | Moderate |
|  | Ni | 0.03 | 0.18 | Low | Moderate |
|  | Pb | 0.09 | 0.57 | Low | Moderate |
|  | Zn | 0.05 | 0.12 | Low | Moderate |
| Adamas | Cd | 0.21 | 0.21 | Moderate | Moderate |
|  | Ni | 0.31 | 0.41 | Moderate | Moderate |
| Paleochori | Cd | 0.04 | **1.84** | Low | **Adverse** |
|  | Ni | 0.71 | **1.01** | Moderate | **Adverse** |
| Methana | Ni | 0.11 | 0.16 | Moderate | Moderate |
|  | Pb | 0.05 | 0.42 | Low | Moderate |

We were especially interested in results from Ischia as it had an abundant *P. oceanica* meadow within the main CO_2_ seep area and has been generally considered a low toxic environment. This seep has the highest Cu (114.60 ± 0.64 mg/Kg) and Pb (14.25±0.29 mg/Kg) concentrations of all the seep locations we sampled, but seagrass tissue analyses showed low levels of these metals (Figs. 3 and 4). On the other hand, the plant compartments of *P. oceanica* at the seeps of Ischia showed higher concentrations of Hg (0.72±0.01 mg/Kg), Ni (5.78± 0.16 mg/Kg), Cd (4.75± 0.60 mg/Kg) and Zn (115.44±1.80 mg/Kg) than the sediment (Fig.3). Interestingly, at the Paleochori seep, Ni (43.27± 2.25 mg/Kg) and Cd (7.76±0.30 mg/Kg) concentrations were very high in the sediments but not within *C. nodosa* plant compartments(Fig.4). Mercury and Zn were also found in high concentrations in the leaves of both *P. oceanica* and *C. nodosa* (Figs. 3 and 4).

**Fig. 4.**
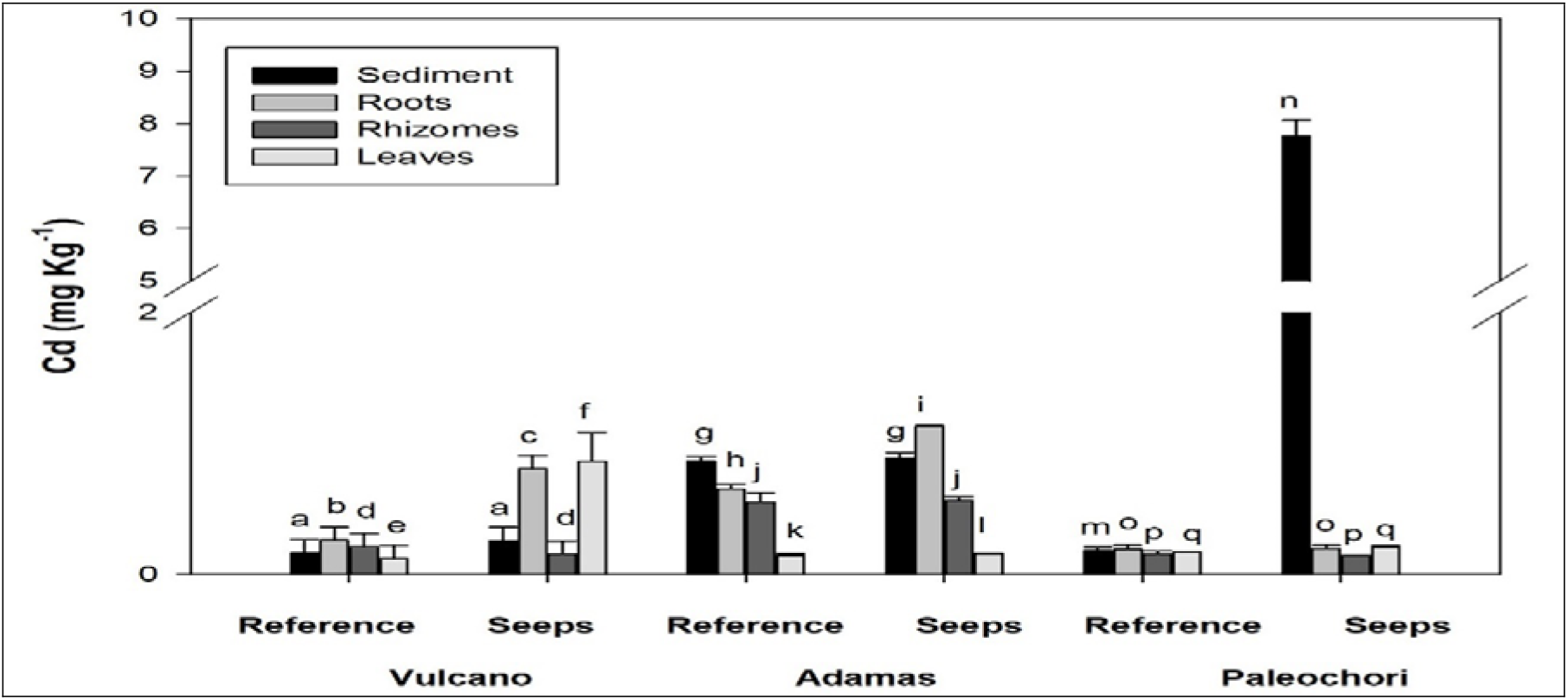
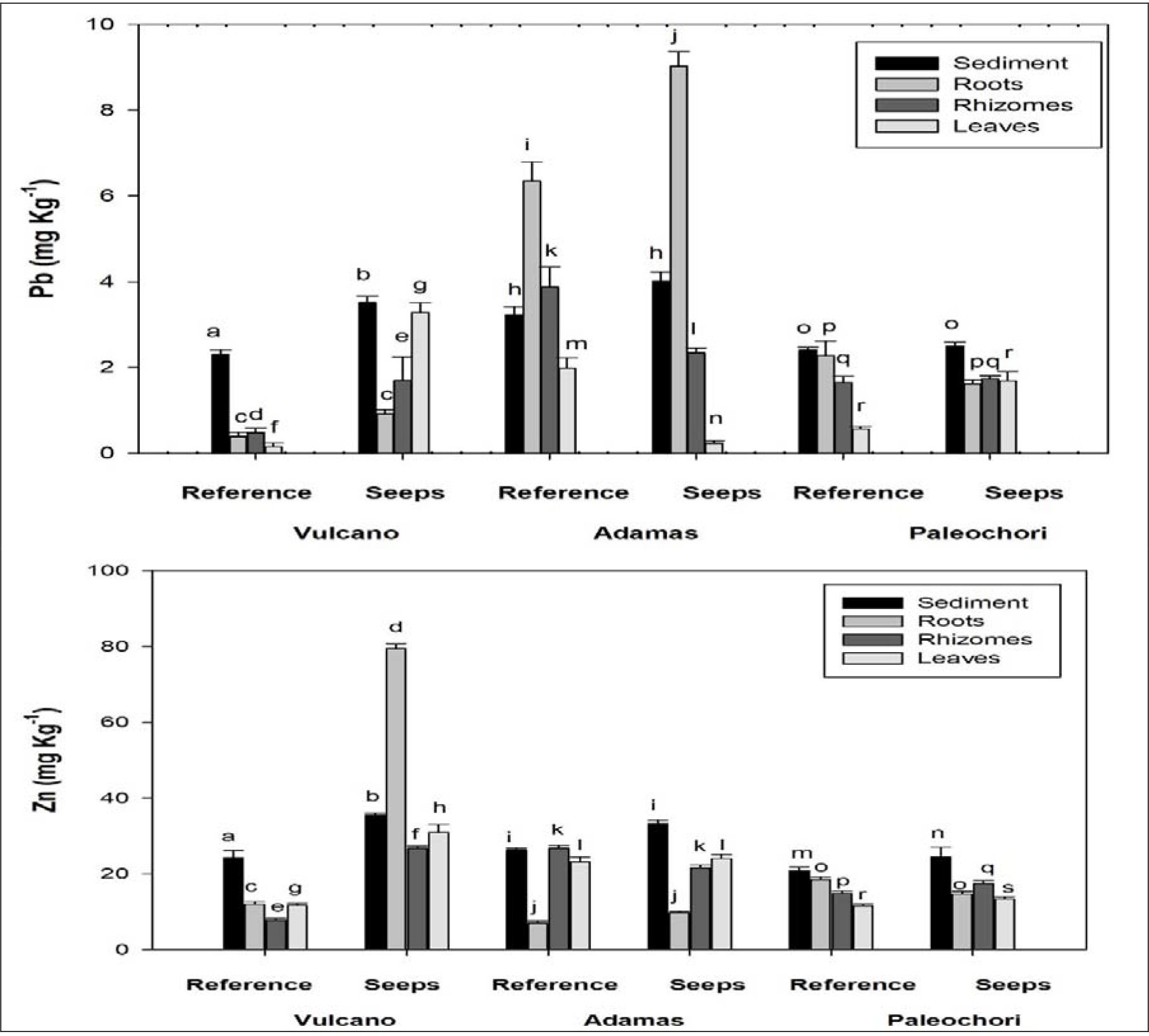
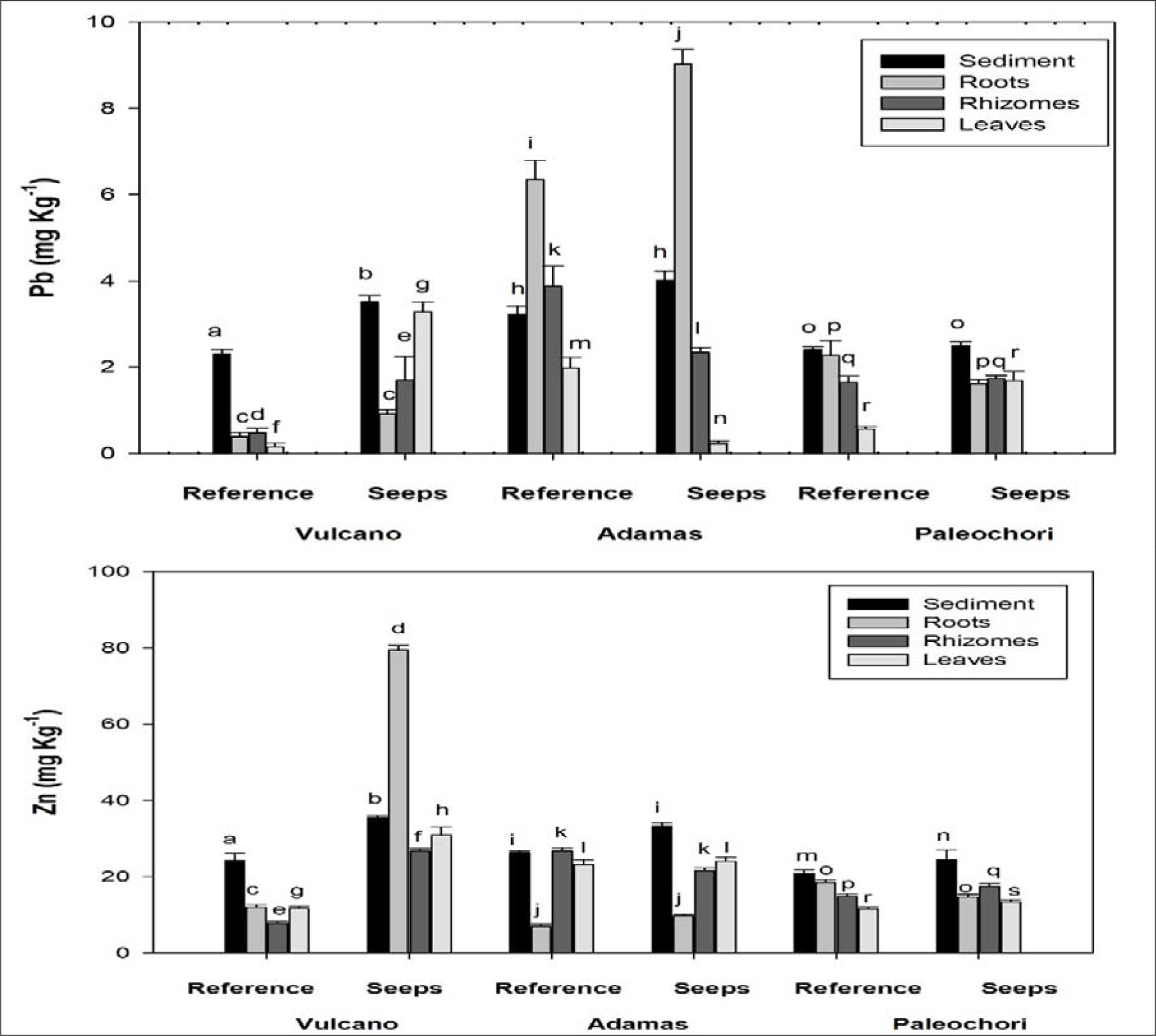
Element concentration (mean ± SE, n=5) of Cd, Cu, Hg, Ni, Pb and Zn for *Cymodocea nodosa* in plant compartments and sediments at reference and CO_2_ seeps off Italy and Greece. Different letters indicate significant differences between reference and CO_2_ seep sites for each location.

Trace element levels were generally significantly higher in the roots than rhizomes and leaves of *P. oceanica* and *C. nodosa* at all seep locations. (Figs.3 and 4) indicating uptake from sediment and storage within plant compartments. Exceptions were the highest Cd (4.75± 0.20 mg/Kg) concentrations within the rhizomes and Cu (58.38±1.01 mg/Kg) within leaves of *P. oceanica* which were not observed in *C. nodosa*. Mercury and Zn were also observed in high concentrations in the leaves of both *P. oceanica* (0.41±0.09; 390.33± 13.48 mg/Kg) and *C. nodosa (0.54*± 0.10; 30.93±2.15 mg/Kg) (Figs.3 and 4).

Significant difference in levels of trace elements in sediment and seagrass compartments were observed for *P. oceanica* between three sites (Table 3). Element concentrations measured in sediments and *P. oceanica* compartments differed significantly except for Cu (sediment-leaves) and Zn (sediment-roots), whereas within *P. oceanica* compartments all elements except Pb (roots-leaves) showed significant difference at all three sites.

**Table 3.**
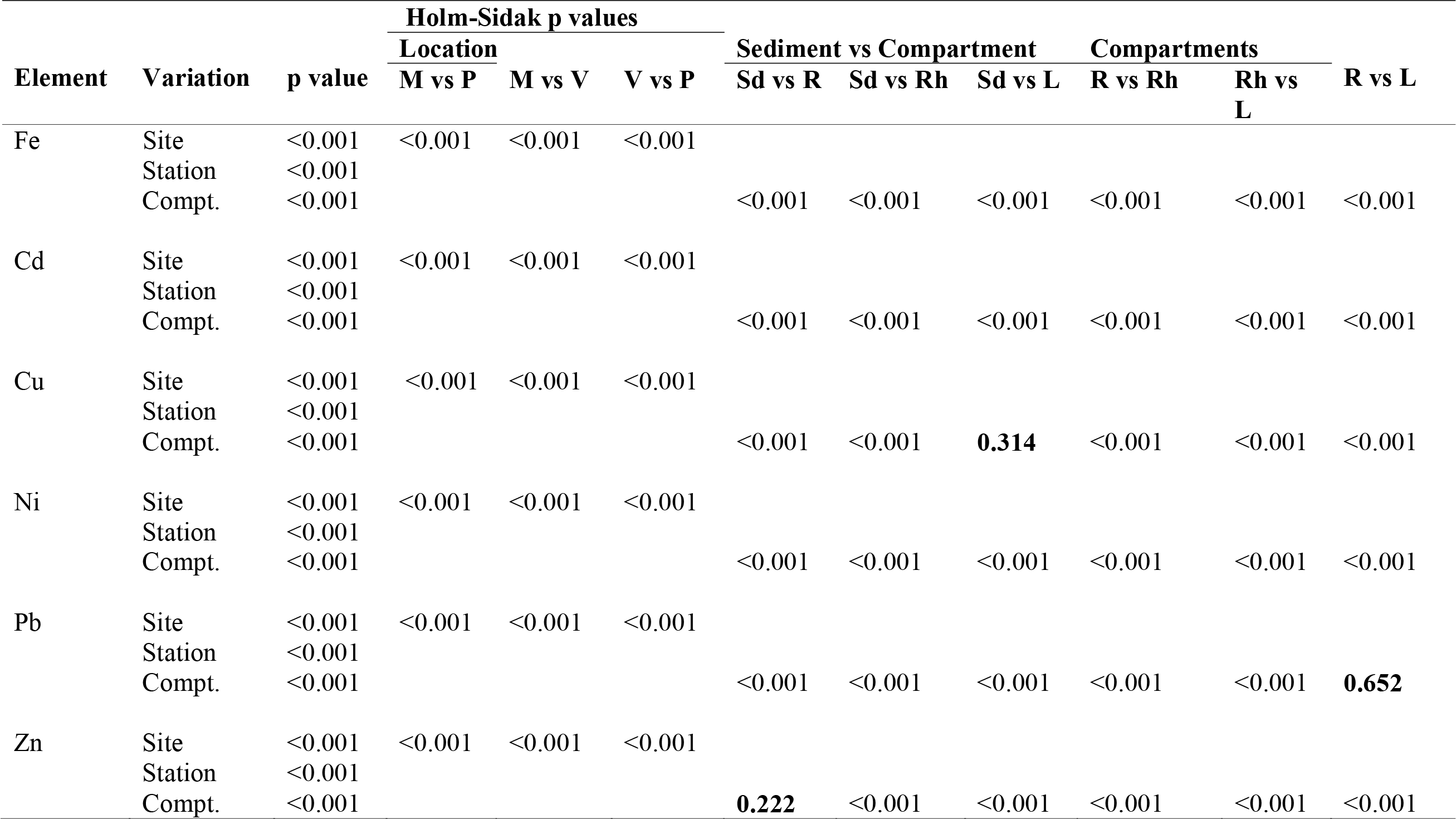
Three-way ANOVA differences in Fe and trace element levels between Location: 3 levels (Methana (M), Panarea(P) and Ischia (V)), Site:2 variables (CO_2_ seeps, Reference)) and compartments :4 levels (Sediments (Sd), Rhizomes (Rh), Roots (R), Leaves (L)). Holm-Sidak significant test (p<0.05) is presented for locations, sediment and *P. oceanica* compartments. Numbers (in bold) indicate differences that were not significant.

| Element | Variation | p value | Holm-Sidak p values |  |  |  |  |  |  |  |  |
| --- | --- | --- | --- | --- | --- | --- | --- | --- | --- | --- | --- |
|  |  |  | Location |  |  | Sediment vs Compartment |  |  | Compartments |  |  |
|  |  |  | M vs P | M vs V | V vs P | Sd vs R | Sd vs Rh | Sd vs L | R vs Rh | Rh vs L | R vs L |
| Fe | Site | <0.001 | <0.001 | <0.001 | <0.001 |  |  |  |  |  |  |
|  | Station | <0.001 |  |  |  |  |  |  |  |  |  |
|  | Compt. | <0.001 |  |  |  | <0.001 | <0.001 | <0.001 | <0.001 | <0.001 | <0.001 |
| Cd | Site | <0.001 | <0.001 | <0.001 | <0.001 |  |  |  |  |  |  |
|  | Station | <0.001 |  |  |  |  |  |  |  |  |  |
|  | Compt. | <0.001 |  |  |  | <0.001 | <0.001 | <0.001 | <0.001 | <0.001 | <0.001 |
| Cu | Site | <0.001 | <0.001 | <0.001 | <0.001 |  |  |  |  |  |  |
|  | Station | <0.001 |  |  |  |  |  |  |  |  |  |
|  | Compt. | <0.001 |  |  |  | <0.001 | <0.001 | **0.314** | <0.001 | <0.001 | <0.001 |
| Ni | Site | <0.001 | <0.001 | <0.001 | <0.001 |  |  |  |  |  |  |
|  | Station | <0.001 |  |  |  |  |  |  |  |  |  |
|  | Compt. | <0.001 |  |  |  | <0.001 | <0.001 | <0.001 | <0.001 | <0.001 | <0.001 |
| Pb | Site | <0.001 | <0.001 | <0.001 | <0.001 |  |  |  |  |  |  |
|  | Station | <0.001 |  |  |  |  |  |  |  |  |  |
|  | Compt. | <0.001 |  |  |  | <0.001 | <0.001 | <0.001 | <0.001 | <0.001 | **0.652** |
| Zn | Site | <0.001 | <0.001 | <0.001 | <0.001 |  |  |  |  |  |  |
|  | Station | <0.001 |  |  |  |  |  |  |  |  |  |
|  | Compt. | <0.001 |  |  |  | **0.222** | <0.001 | <0.001 | <0.001 | <0.001 | <0.001 |

Significant variation was observed in trace elements level for *C. nodosa* between the three sites except for Cu at Adamas vs Paleochori and Ni at Vulcano vs Adamas (Table 4). Element levels measured in sediment and in *C. nodosa* compartments differed significantly except for Cu (sediment vs rhizomes), whereas within plant compartments significant difference was not observed for Zn (rhizomes vs leaves) and Cd (rhizomes vs leaves) at all three sites.

**Table 4.**
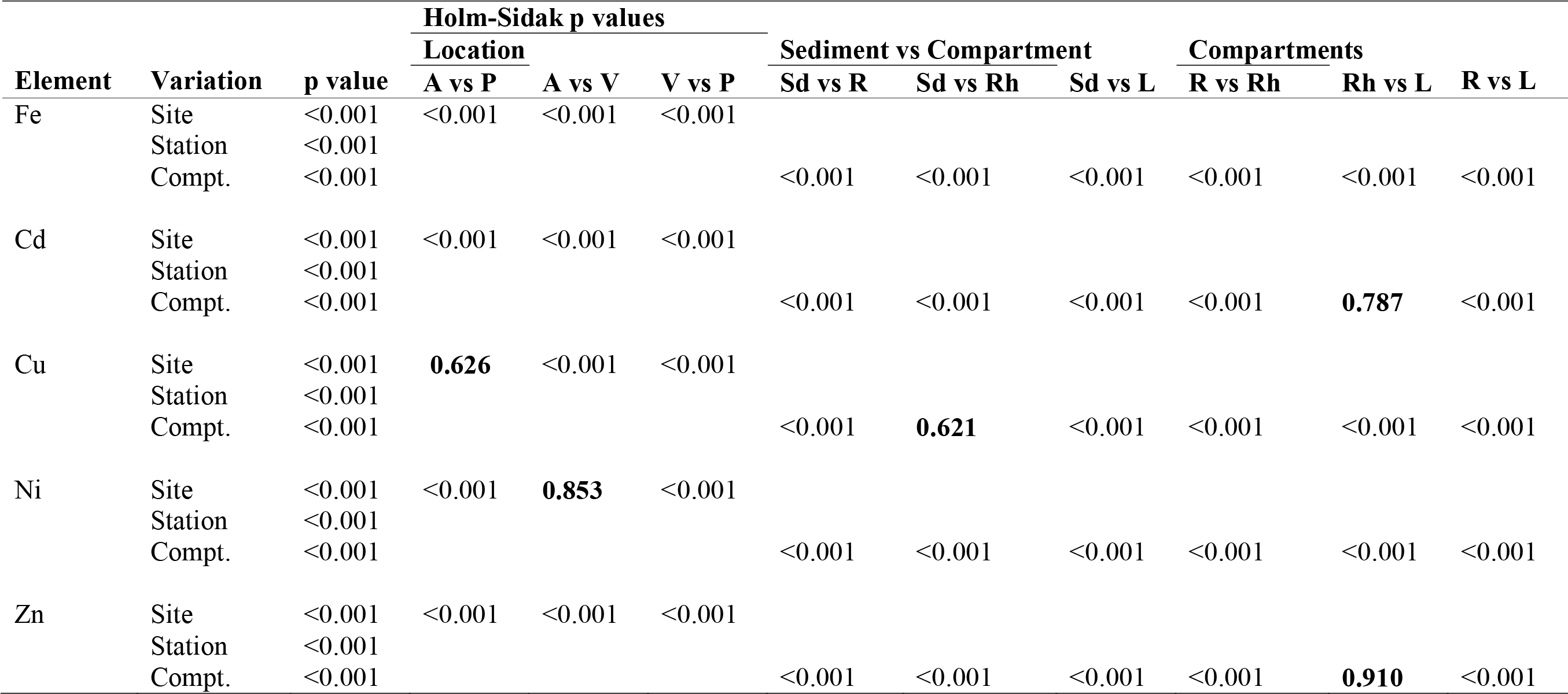
Three-way ANOVA differences in Fe and trace element levels between Location: 3 levels (Adamas (A), Paleochori (P) and Vulcano (V)), Sites:2 variables (CO_2_ seeps, reference) and compartments: 4 levels (Sediments (Sd), Rhizomes (Rh), Roots (R), Leaves (L). Holm-Sidak significant test (p<0.05) is presented for locations, sediment and *C. nodosa* compartments. Numbers (in bold) indicate differences that were not significant.

The Bio Sediment Accumulation Factor index indicated that both *C. nodosa* and *P. oceanica* roots and leaves accumulated trace elements from sediment. In *P. oceanica* Cd, Cu, Hg, Ni, Pb and Zn were observed with BSAF>1 in the roots or leaves at all three CO_2_ seeps. Trace elements Cd, Cu, Ni and Zn were observed with BSAF >1 in the roots and leaves of *C. nodosa* at all three CO_2_ seeps. BSAF>1 for elements in both seagrass roots and leaves indicate these parts of the plant accumulated higher level of elements at CO_2_ seeps.

Correlations between trace element content in sediments and those recorded in *P. oceanica* compartments were significant and positive for Zn and Ni in rhizomes at Ischia and Panarea seeps respectively (Table 5). Correlations were negative for Ni and Cd in rhizomes at Methana and Panarea seeps respectively (Table 5). The correlation for element content between any two organs of *P. oceanica* were positive and significant for Cd and negative for Cu at Ischia CO_2_ seeps (Table 5).

**Table 5.** Correlation between trace elements in sediments versus *P. oceanica* roots, rhizomes and leaves and plant compartments at Mediterranean CO_2_ seeps. The correlation co-efficient (r) and significance level (p) are presented. Numbers in bold indicate significant correlation, only trace elements with significant correlations are shown.

| Location | Element | Sediment-roots |  | Sediment-rhizomes |  | Sediment-Leaves |  |
| --- | --- | --- | --- | --- | --- | --- | --- |
|  |  | r | p | r | p | r | p |
| Ischia | Zn | -0.234 | 0.704 | 0.870 | **0.048** | 0.321 | 0.598 |
| Panarea | Cd | 0.841 | **0.014** | -0.910 | **0.032** | -0.064 | 0.918 |
|  | Ni | -0.358 | 0.554 | 0.884 | **0.046** | -0.787 | 0.114 |
| Seagrass compartments |  |  |  |  |  |  |  |
| Location | Element | Roots-Rhizomes |  | Roots-Leaves |  | Rhizomes-Leaves |  |
|  |  | r | p | r | p | r | p |
| Ischia | Cd | 0.300 | 0.683 | 0.975 | **0.016** | 0.359 | 0.517 |
|  | Cu | -0.273 | 0.657 | 0.877 | **0.049** | -0.577 | 0.308 |

Correlations of trace element content in sediment and those found in rhizomes of *C. nodosa* at Vulcano were significant and positive, whereas significant and negative correlation were observed for Zn content between sediment and rhizomes and leaves (Table 6). For any two plant organs Cd was found with positive co-relation at Vulcano, where significant and negative correlation was observed for Cu and Hg at Vulcano and Cu at Adamas (Table 6).

**Table 6.** Correlation between trace elements in sediment versus C. *nodosa* roots, rhizomes and leaves and between plant compartments at Mediterranean CO_2_ seeps. The correlation co-efficient (r) and significance (p) level are presented. Numbers in bold indicate significant correlations, only trace elements with significant co-relation are shown.

| Location | Element | Sediment-roots |  | Sediment-rhizomes |  | Sediment-Leaves |  |
| --- | --- | --- | --- | --- | --- | --- | --- |
|  |  | r | p | r | p | r | p |
| Vulcano | Fe | 0.437 | 0.462 | 0.992 | **0.000** | -0.836 | 0.078 |
|  | Zn | -0.795 | 0.108 | -0.966 | **0.007** | -0.906 | **0.034** |
| Seagrass Compartments |  |  |  |  |  |  |  |
| Location | Element | Roots-Rhizomes |  | Rhizomes-Leaves |  | Roots-Leaves |  |
|  |  | r | p | r | p | r | p |
| Vulcano | Cd | 1.000 | **0.016** | 0.158 | 0.783 | 0.158 | 0.783 |
|  | Cu | -0.986 | **0.002** | 0.534 | 0.354 | -0.620 | 0.265 |
|  | Hg | -0.135 | 0.783 | 0.216 | 0.683 | -0.947 | **0.016** |
| Adamas | Fe | -0.975 | **0.016** | -0.300 | 0.683 | 0.205 | 0.683 |
|  | Cu | -0.872 | 0.083 | -1.000 | **0.016** | 0.872 | 0.083 |

## Discussion

Shallow water CO_2_ seeps have been used as natural analogues for future coastal ecosystems as they can have areas of seabed where entire communities of marine organisms are exposed to the shifts in carbonate chemistry that are expected due to continued anthropogenic CO_2_ emissions (Hall-Spencer et al., 2008; Enochs et al., 2015; Connell et al., 2017). At such seeps, there are often elevated levels of H_2_S and trace elements (Vizzini et al., 2010; Kadar et al., 2012; Boatta et al., 2013) so care is needed when using them to assess the effects of low pH due to confounding factors that may be harmful to marine biota (Bary et al., 2010). The six CO_2_ seeps that we survey had gradients in seawater pH conditions with sediments that were enriched with Cd, Cu, Hg, Ni, Pb and Zn. This was expected since hydrothermal seeps sediments often have high levels of metals (Aiuppa et al., 2000; Sternbeck et al., 2001) due to continuous input from the subsea floor into the sediments (Dando et al., 2000; Hall-Spencer et al., 2008). Our calculated Sediment Quality Guidelines Quotient (Long et al., 1998; MacDonald et al., 2000) suggests Hg (at Vulcano), Cu (at Ischia) plus Cd and Ni (at Paleochori) were at high enough levels to possibly have adverse impacts on seagrass associated biota.

The mean trace element range measured within the CO_2_ seep sediments in our research were lower compared to mean element levels observed around Mediterranean coast of Italy and Greek (Table 7), except for Cu (12.10-114.60) at Italy and Ni (6.91-44.37) at Greek sites (Figs 3 and 4). However, trace element levels observed at the sediments of CO_2_ seeps of Vulcano, Italy were in the same range measured by Vizzini et al. (2013), whereas for sediments of Panarea CO_2_ seeps, Pb concentration in our results (Fig. 3) were 5-fold lower from the findings of Monia Renzi et al. (2011). Lower levels of trace elements in the sediments can be due to lower level of input from CO_2_ seeps compared to the anthropogenic input the Mediterranean coast receives and the sediment redox potential values which determines the binding and deposition of elements in the sediments (Monia Renzi et al. 2011). Secondly, the binding of trace elements in the sediments depends on the clay particles (<63μm), which were significantly lower in our studies as majority of the sediments were sand at CO_2_ seeps.

**Table 7.** Mean range of trace element (mg/Kg) levels measured in surface sediments and in *P. oceanica* and *C. nodosa* compartments (roots, rhizomes and leaves) off Italy and Greek coast in Mediterranean Sea.

| Sample | Location | Tissues | Cd | Cu | Hg | Ni | Pb | Zn |
| --- | --- | --- | --- | --- | --- | --- | --- | --- |
| Sediment | Italy |  | 0.21-1.92 | 14.92-55.52 | 0.61-2.57 | 13.26-201.45 | 23.67-45.46 | 49.48-76.81 |
|  | Greek |  | 0.22-43.57 | 30.64-204 | 0.05-0.39 | 42.20 | 55.42-309 | 128-225.60 |
| *P. oceanica* | Italy | Leaf | 1.17-3.16 | 6.97-16.17 | 0.06-0.11 | 16.01-55.08 | 2.25-6.43 | 113.02-203.38 |
|  |  | Rhizome | 0.55-1.51 | 8.37-21.37 | 0.08-0.11 | 3.82-8.29 | 3.19-10.50 | 54.41-171 |
|  |  | Root | 0.70-1.23 | 13.33-17.40 | 0.21-0.34 | 6.20-8.56 | 4.94-9.63 | 65.86-119.67 |
|  | Greek | Leaf | 6.52 | 9.66-45-80 | 0.07 | 18.45-60.90 | 15.61 | 88.20 |
|  |  | Rhizome | 0.53 | 2.75-58.6 | - | 13.17-46.20 | 15.20 | 59.88 |
|  |  | Root | 0.74 | 5.37-36.10 | - | 7.27-46.21 | 43.10 | 55.67 |
| *C. nodosa* | Italy | Leaf | 0.28-2.73 | 7.30-28.81 | 0.27-2.23 | 3.84-6.65 | 2.61-23.56 | 48.8-65-8 |
|  |  | Rhizome | 0.05-0.22 | 5.69-23.1 | 0.13 | 1.52-5.34 | 0.45-1.87 | 21.15-31.17 |
|  |  | Root | 0.22-0.45 | 6.57-25-8 | 0.19 | 4.05-7.89 | 4.58-6.99 | 37.25-50.44 |
|  | Greek | Leaf | 0.70-0.79 | 7.75-59.60 | - | 3.40-27.03 | 108-210 | 87.27-627 |
|  |  | Rhizome | 0.89-0.44 | 4.95-17.65 | - | 0.90-6.11 | 117-220 | 45.47-77.70 |
|  |  | Root | 0.89-0.44 | 7.58-49.80 | - | 2.28-19.43 | 113.50-220 | 34-77 |

Element levels were higher in seagrass compartments at the seep sites compared to reference sites. Mean range of elements measured in *Posidonia oceanica* leaves, rhizomes and roots at CO_2_ seeps were higher, lower or equivalent to mean range of elements observed around Mediterranean coast of Greek and Italy (Table 7). Elements such as Cu (2.4-fold) and Hg (4-fold) in leaves, Cd (1.6-fold) and Hg (1.4-fold) in rhizomes and Cu (2.1-fold) and Hg (1.6-fold) in roots at CO_2_ seeps of Ischia and Panarea were higher than the levels observed for *P. oceanica* off Italy coast (Table 7). However, the element levels at the CO_2_ seeps of Methana for *P. oceanica* compartments were lower than the element levels observed off Greek coast, except for Ni in rhizomes (Table 7). *C. nodosa* element levels at CO_2_ seeps of Milos Islands were lower compared to element levels of Greek coast (Table 7) except for Ni (4-fold) in rhizomes and Cd (1.5-fold) and Cu (2-fold) in roots, which were higher. However, at the seeps of Vulcano elements such as Cd (2-fold), Cu (9.8-fold), Hg (2-fold), Ni (4.8-fold) and Zn (1.6-fold) in roots, Cu (1.1-fold) and Hg (1.3-fold) in rhizomes and Ni (1.6-fold) in leaves were higher than element levels observed for Italy coast (Table 7). This suggests that, there is multi-fold variation in element input and accumulation from the sediments at the CO_2_ seeps at different sites and species, which has been observed for *P. oceanica* and *C. nodosa* around the Mediterranean coast (Bonanno and Bonaca,2017). This also agrees with the fact that seagrass element accumulation is more element and seagrass tissue-specific rather than species-specific (Bonanno and Bonaca, 2017) resulting in seagrass compartments acting as metal accumulators of their surrounding environment, especially of heavy metals (Govers et al., 2014).

Majority of accumulation of elements in both seagrasses at CO_2_ seeps were observed in roots >rhizomes > leaves, which is common for *P. oceanica* and *C. nodosa* (Bonanno and Bonaca,2017). Root accumulation is common in both terrestrial and aquatic plants where they store certain elements to avoid damage to photosynthetic apparatus. However, significant population declines due to higher element levels in root biomass have not been reported (Lafabrie et al., 2007). Saying that, the storage and translocation of elements such as Cd, Ni and Pb as observed at CO_2_ seeps within seagrass compartments from roots-rhizomes- leaves, suggests seagrass may adopt a strategy depending on their physiology to either accumulate elements in the below ground root biomass or move out the elements through the leaves which are then shed, as observed in *P. oceanica* (Di Leo et al., 2013; Richir and Gobert, 2016) and in *C. nodosa* (Malea and Haritinoids, 1999; Bonanno and Di Martino, 2016). Other findings have also reported that seagrasses prefer to accumulate certain elements such as Cd and Ni that are essential micronutrients (Sanz-Lazaro et al., 2012) rather than Hg or Pb that are toxic (Kabata-Pendias and Mukherjee, 2007), which has been observed for accumulation of Zn over Pb in both *P. oceanica* (Sanchiz et al., 2001) and *C. nodosa* (Malea and Haritonidis, 1999; Llagostera et al., 2011).

Correlation results between elements in sediment and that in seagrass compartments indicated that the preferable accumulation pattern of elements from sediments are not always the sediment-root pathways, even though higher element concentrations were observed in the sediments at CO_2_ seeps. For instance, in *P. oceanica* Cd, was found with a negative correlation through sediment-root and Cu, Ni and Zn through root-rhizome-leaves pathway, whereas in *C. nodosa* negative correlation was found for Zn between sediment-rhizomes and leaves. Negative correlation suggests that the preferable route for Cd, Cu, Ni and Zn transfer and translocation in *P. oceanica* (Lafabrie et al., 2007; Di Leo et al., 2013) and Zn in *C. nodosa* (Malea et al., 1999) compartments is through water column at CO_2_ seeps rather than the sediment-root pathways. Similarly, elements such as Hg with negative correlation in *C. nodosa*, suggests Hg being toxic is not allowed for translocation within the seagrass compartments (Sanchiz et al., 2001). Similar results of transfer and translocation of elements (both essential and toxic) within seagrass compartments were observed for *P. oceanica*, *C. nodosa* and *Halophila stipulacea* of Mediterranean Sea, where the seagrass species have shown high element mobility from water column and highly variable element and species-specific translocation capabilities (Malea and Kevrekidis, 2013; Bonanno et al., 2017; Bonanno and Raccuia, 2018).

Correlation data between both seagrass compartments are different because element accumulation patterns in seagrass are governed by multitudes of factors (Llagostera et al., 2011). Higher element concentrations in seagrass roots, rhizomes and leaves in our studies at CO_2_ seeps indicate the capacity of seagrass to absorb element simultaneously from sediments and water, as they are always submerged. Absorbing elements simultaneously at these CO_2_ seeps have helped the seagrasses to adapt and manage stressful element levels found in sediments and regulate their gene expression to increased metal stress which have been observed for *P. oceanica* by Lauritano et al., (2015) and for *C. nodosa* by Olive et al., (2017).

At CO_2_ seeps the low pH can alter the metal speciation and favour the release of metals from sediment (Simpson et al., 2004: Atkinson et al., 2007). The chemical form in which metals are present (e.g. whether they are bound to organic or inorganic compounds) is a key issue determining its bioavailability. Low pH of seawater near the CO_2_ seeps tends to release the metals that are less strongly associated with sediments, increasing their potential bioavailability (Riba et al., 2004). Thus, low pH can increase the concentration of certain dissolved metals, which could affect the sediment-seagrass associated biota e.g., by increasing Cu, Cd and Zn bio-availability, their accumulation and possible toxic effects (Basallote et al., 2014).

In our research, all the CO_2_ seeps had low pH (7.4-7.9) conditions, which are known to increase the availability of Cd, Cu, Ni, Pb and Zn in their free ion forms (Roberts et al., 2013). Low pH combined with increased availability can influence and increase seagrass uptake of trace elements (Yang and Ye, 2009) that can lead to higher accumulation and storage of trace elements in seagrass roots and leaves (Bonanno and Bonaca, 2017). Higher accumulation can lead to metal stress once threshold levels are reached and affect the seagrass physiological processes (Olive et al., 2017). However, it is difficult to measure toxic effects of metals on seagrass in *in-situ* conditions due to variable environmental settings, but a few *ex-situ* studies on metal toxicity have been conducted on *Cymodocea serrulata* (Prange and Dennison. 2000), *Halophila ovalis* and *H. spinulosa* (Prange and Dennison, 2000; Ambo-Rappe et al., 2011). Considering the observed results from these ex-situ metal toxicity studies, there is a possibility that elements such as Cu and Pb concentrations at the CO_2_ seeps may affect *P. oceanica* and *C. nodosa* photosynthesis as well as root and leaf structures (Prange and Dennison. 2000; Ambo -Rapee et al., 2011). This can be one of the reasons due to which seagrasses are abundant at some seeps (e.g. Ischia) but not others (e.g. Vulcano) given that they grow well and can take advantage of elevated CO_2_ levels at some seeps but not at others (Vizzini et al., 2010; Russell et al., 2013; Olive et al., 2017).

## Conclusion

To sum up: we observed that Mediterranean CO_2_ seep sites of Greek and Italy consistently have elevated levels of trace elements in sediments, which can be used to study the interactions between low pH, element bioavailability and element accumulation within seagrasses compartments. We concur with Bary et al., (2011), Vizzini et al., (2013), Lauritano et al., (2015) and Olive et al., (2017) that care is needed when using volcanic CO_2_ seeps as analogues for the effects of ocean acidification as there can be areas with increased levels of elements that can be harmful to seagrasses. In some cases, such as Ischia, high levels of trace elements available in the sediment, such as Cu were not accumulated in seagrass, whereas in Vulcano, we found that elevated levels of Zn were accumulated in seagrass roots and rhizomes. Thus, future work that use seep sites to assess the effects of trace element availability and accumulation in seagrass ecosystems can shed light on trace element impacts on seagrass physiology, threshold levels of trace elements and their toxic effects on seagrass and probably define sediment quality guidelines for seagrass and seagrass associated biota (Bouchon et al., 2016) under low pH conditions. The findings of our research are relevant to agencies responsible for monitoring the effects of trace element contamination in the marine environment using seagrass ecosystems, conservation and protection of seagrass ecosystems in the Mediterranean Sea for their ecosystem services.

## Acknowledgement

This work was part of MARES “Future Oceans” project (MARES _12_14) and was funded through a MARES Grant. MARES is a Joint Doctorate programme selected under Erasmus Mundus coordinated by Ghent University (FPA 2011-0016). Check http://www.mares-eu.org for extra information. We are grateful to Dr Marco Milazzo for his support during the field work at Vulcano, Italy. Dr Joao Silva and Dr Irene Oliva for helping me collect samples form Ischia and Panarea, Italy. We are grateful for the support of Thanos Dailianis, Julius Glampedakis in collection of samples from Greece and Dr Eugenia Apostolaki for her support during the field work. I am thankful to Andrew Tonkin and Robert Clough at Plymouth University, UK for helping me in the laboratory analysis. We would like to thank Prof. Paul Dando and Prof. Franceso Parello for their constructive comments on an early draft.

